# Individual differences in representational similarity of first and second languages in the bilingual brain

**DOI:** 10.1101/2021.01.18.427141

**Authors:** Emily S. Nichols, Yue Gao, Sofia Fregni, Li Liu, Marc F. Joanisse

## Abstract

Current theories of bilingualism disagree on the extent to which separate brain regions are used to maintain or process one’s first and second language. The present study took a novel multivariate approach to address this question. We examined whether bilinguals maintain distinct neural representations of two languages; specifically, we tested whether brain areas that are involved in processing word meaning in either language are reliably representing each language differently, and whether language representation is influenced by individual differences in proficiency level and age of acquisition of L2. Thirty-one English-Mandarin bilingual adults performed a picture-word matching task in both languages. We then used representational similarity analysis to examine which brain regions reliably showed different patterns of activity for each language. As a group, there were no regions that reliably represented languages distinctly. However, both proficiency and age of acquisition predicted dissimilarity between language representations in several brain areas within the language network as well as several regions of the ventral visual pathway, demonstrating that top-down language knowledge and individual language experience shapes concept representation in the processing stream. The results support the model of an integrated language system in bilinguals, along with a novel description of how representations for each language change with proficiency level and L2 age of acquisition.

## 1. Introduction

Current theories of bilingual language processing hold that bilingual speakers coactivate their two languages during speech, and that they maintain similar, overlapping representations for both ^1–4^. Additionally, past neuroimaging research has provided much evidence that a second language (L2) is processed similarly to the speaker’s first language ^5–9^. Even when L1-L2 differences exist, such as more extensive activity in L2 ^10,11^, there remains extensive overlap ^6^. This suggests that similar underlying language networks are engaged regardless of which language is being used. The concept of language coactivation in bilinguals is widely accepted, as is that of a single, integrated lexicon ^12,13^. While neuroimaging provides much support for an integrated lexicon through activation of similar structures, the separation of patterns of activity within the shared L1/L2 brain areas may provide evidence for some degree of distinction between L1 and L2 lexicons.

Despite L1 and L2 sharing a network of structures, traditional univariate contrasts cannot tell us how languages are being represented in those areas, and while there is extensive overlap in brain areas that represent L1 and L2 ^6,7,9,14^, how the languages are represented may vary. That is, regions coding for language-specific information, such as spoken codes (e.g., left superior temporal gyrus and left inferior frontal gyrus) may represent each language differently. In contrast, regions involved in executive and attentional control (e.g., dorsolateral prefrontal cortex and insula) are likely to show less differentiation in how each language is represented as the function of these regions should not differ qualitatively from one language to another. Individual differences in language ability and experience also play an important role in bilingual language processing ^11,15,16^, and may affect the integration of the neural representation of each language. Previous research indicates that low proficiency speakers and late L2 learners have greater separation of their two languages’ conceptual knowledge ^17^, and this separation may also be reflected in the neural representation of words and concepts within co-activated brain areas.

Representational Similarity Analysis (RSA) is an fMRI analysis technique relying on reproducible spatial patterns of activity that correlate with distinct experimental conditions ^18^, and has been used in the past to identify regions that differentiate between languages during reading ^19^. RSA has been used to reveal differences between conditions within individual brain regions that were previously undetectable using standard univariate methods; it reveals cortical patterns sensitive to differences in stimuli even when the degree of activation is similar ^20–23^. This technique may be particularly relevant to describing bilingual word processing, as it has the potential to identify differences between languages that were previously thought to not exist. RSA allows us to examine possible language-processing differences in areas that are assumed to be engaged similarly for both languages, suggesting that they are representing L1 and L2 differently. Additionally, by measuring individual differences in bilingual experience, it is possible to determine how the neural representation of each language changes with these measures.

### Rationale for the Present Study

The present study examined whether brain areas involved in both L1 and L2 representationally distinguish the two languages. English-Mandarin bilingual adults performed a lexico-semantic recognition task in each language. We then examined whether brain regions showed reliably different patterns of activity for each language within regions that significantly activated to both. We predicted that, consistent with models of an integrated bilingual lexicon, representational dissimilarity would decrease with increasing proficiency and earlier ages of L2 acquisition (AoA). In contrast, areas involved in domain general cognitive processes, such as executive function, were not expected to show language-selective patterns.

## 2. Material and methods

### 2.1 Participants

Thirty-two (13 female) neurologically healthy right-handed native speakers of English were recruited via posters and word of mouth in Beijing, China. All participants were second-language learners of Mandarin, aged 18-37 (*M* = 23.84, *SD* = 4.59), and had begun learning Mandarin between the ages of 0-28 years (*M* = 18.09, *SD* = 7.10). This study was approved by the Beijing Normal University research ethics board and all participants gave informed consent prior to participation. Demographic and language information is summarized in Table 1.

**Table 1.** Participant demographic and language information

| Measure | M (SD) |
| --- | --- |
| N | 32 |
| Sex | 13 female, 19 male |
| Age (years) | 23.84 (4.59) |
| Age of L2 acquisition | 18.90 (7.10) |
| *Proficiency (%)* |  |
| English | 88.93 (5.88) |
| Mandarin | 38.54 (18.15) |
| *Reaction Time (ms)* |  |
| English | 1203.60 (202.21) |
| Mandarin | 1607.88 (231.78) |
| *Accuracy (%)* |  |
| English | 94.17 (4.21) |
| Mandarin | 83.07 (10.35) |

### 2.2 Behavioral tests

L1 English and L2 Mandarin proficiency levels were assessed prior to scanning using a subset of 48 questions from the Test of English as a Second Language (ETS, Princeton, NJ) and 48 questions from the Hanyu Shuiping Kaoshi (HSK Centre, Beijing, China), respectively. Both tests consisted of three sections, grammar, reading comprehension, and vocabulary, which were combined to give a final score for each language, representing overall proficiency in these three domains.

Age of acquisition was obtained by self-report, defined as the age at which individuals first began learning Mandarin. To verify handedness, participants completed an abridged version of the Edinburgh Handedness Inventory ^24^. Behavioral measures, informed consent and task instructions were administered in English, aside from the Mandarin proficiency test, which was administered in Mandarin.

### 2.3 fMRI Task

Participants completed a picture-word matching task during scanning, in alternating runs of English and Mandarin. Pictures were presented via LCD projector to the center of a screen mounted at the head of the scanner bore, which was viewed through a mirror placed above the head coil. At the same time, a word was played binaurally through insert earphones (Sensimetrics Corporation, Malden, MA). Participants were required to indicate as quickly as possible with a button press whether the picture and word matched. Each picture was visible for 2.5 s. They viewed a fixation crosshair between trials as baseline. Stimulus presentation and response recording was controlled with E-Prime software (Psychology Software Tools, Inc., Sharpsburg, PA) and a Windows laptop.

The scanning session was divided into 8 alternating English and Mandarin runs. Four English runs were interleaved with four Mandarin runs, with starting language counterbalanced, so that a run in the first language was always followed by a run in the other language. Four orderings were produced: one version starting with English, one version starting with Mandarin, and an additional version of each in which runs were presented in the reverse order. Each run began with an image reminding participants of which buttons to respond with, and the language in which the next run would be performed. Each run consisted of 20 trials for a total of 160 trials (80 in each language, with 40 matching and 40 mismatching). A short break was provided between each 3.5-minute scanning run. Each image appeared twice during the experiment, once in a matching pair and once in a semantically unrelated mismatching pair. Each trial was 2.5 s in duration, with inter-trial interval jittered between 2.5 and 12.5 s in 2.5 s increments, to optimize the deconvolution of the blood oxygen level dependent signal.

Stimulus words consisted of 40 common single-word concepts with the constraint that they are expressed as single two-syllable words in both English and Mandarin, and have frequencies greater than 40 per million in both languages (English: CELEX Lexical Database ^25^ and Mandarin: SUBTLEX-CH ^26^). In a separate pilot study involving different participants, we asked groups of native speakers of English or Mandarin to rate the imageability and familiarity of the stimulus words, as well as the correspondence of the pictures to target words, on a Likert scale of 1-7. Both groups showed equally high ratings on familiarity (*M*_*Mandarin*_ = 5.78, *M*_*English*_ = 5.48) and picture/word correspondence (*M*_*Mandarin*_ = 6.08, *M*_*English*_ = 5.95).

### 2.4 Data acquisition and processing

Imaging was conducted on a Siemens Magnetom TIM Trio whole-body 3 Tesla scanner with a 32-channel head coil. T2*-weighted functional scans were acquired in the transverse plane with 45 slices per volume (TR = 2.5 s; TE = 38 ms; flip angle = 80°; FOV = 192 × 192 mm; voxel size 3×3×3 mm^3^) using an iPAT parallel acquisition sequence (generalized auto-calibrating partially parallel acquisition [GRAPPA]; acceleration factor = 2), providing full coverage of the cerebrum and the superior portion of the cerebellum. A total of 576 functional scans were acquired for each participant over 8 runs (3.5 min per run). After the final functional run, a whole-head high-resolution 3D anatomical scan was acquired in the sagittal plane, using a 3D pulse sequence weighted for T1 contrast (MPRAGE; TR = 2.3 s; TE = 2.98 ms; FOV = 256 × 256 mm; voxel size = 1 mm^3^; 176 slices; GRAPPA acceleration factor = 2).

Raw data were converted from DICOM to BIDS format and preprocessed using FMRIPREP version 1.0.0 ^27^ a Nipype ^27,28^ based tool. Each T1 weighted volume was corrected for bias field using N4BiasFieldCorrection v2.1.0 ^29^ and skullstripped using antsBrainExtraction.sh v2.1.0 (using OASIS template). Cortical surface was estimated using FreeSurfer v6.0.0 ^30^. The skullstripped T1w volume was coregistered to skullstripped ICBM 152 Nonlinear Asymmetrical template version 2009c ^31^ using nonlinear transformation implemented in ANTs v2.1.0 ^32^.

Functional data was slice time corrected using AFNI ^33^ and motion corrected using MCFLIRT v5.0.9 ^34^. This was followed by co-registration to the corresponding T1-weighted volume using boundary based registration 9 degrees of freedom - implemented in FreeSurfer v6.0.0 ^35^. Motion correcting transformations, T1 weighted transformation and MNI template warp were applied in a single step using antsApplyTransformations v2.1.0 with Lanczos interpolation.

Three tissue classes were extracted from T1w images using FSL FAST v5.0.9 ^36^. Voxels from cerebrospinal fluid and white matter were used to create a mask in turn used to extract physiological noise regressors using aCompCor ^37^. Mask was eroded and limited to subcortical regions to limit overlap with gray matter, six principal components were estimated. Frame-wise displacement ^38^ was calculated for each functional run using Nipype implementation. For more details of the pipeline see https://fmriprep.readthedocs.io/en/latest/workflows.html.

### 2.5 First- and second-level statistics

Single-subject statistical maps were formed in the context of the General Linear Model using AFNI 3dDeconvolve function. Linear trends in the functional data were removed, and first-level analysis was conducted by modeling all English trials together and all Mandarin trials together. The statistical maps were formed in the context of the General Linear Model using AFNI 3dDeconvolve function. Additional regressors were included for the six motion parameters, physiological noise from the preprocessing step, and the response times. This led to one English and one Mandarin output per subject that we used to compute the univariate contrasts. One sample t-tests against zero were then computed for each language (AFNI 3dttest++) and a conjunction analysis (AFNI 3dcalc) was performed to identify areas that significantly activated for both English and Mandarin. The result was a conjunction map thresholded at 2.596 (*p* = 0.01 uncorrected); a fairly liberal threshold was used at this stage in order to include as many areas in the search space as possible. A brain mask was then created using the results of this conjunction analysis. Finally, first-level single-subject statistics were recomputed for English and Mandarin, this time creating separate models for even and odd runs. Only correct trials were included in both first-level analyses, with accuracy ranging from 81.25% to 100% correct on the English task and ranging from 61.25% to 96.25% correct on the Mandarin task.

### 2.6 Split-half correlation searchlight analysis

Searchlight RSA was then performed to identify regions in which the representations of L1 and L2 were reliably different, regardless of groupwise differences in activation levels. The search space for the analysis was constrained to regions within the English-Mandarin conjunction mask, shown in Figure 1. To conduct RSA, a split-half correlation searchlight was performed within the CoSMoMVPA Matlab toolbox ^39^, using a search sphere radius of 3 voxels. Within each searchlight sphere Pearson correlations were performed for activity patterns between even and odd runs, within-language (English-English and Mandarin-Mandarin) and between-language (English-Mandarin), yielding a 2 × 2 similarity matrix for each individual at each point of the searchlight. Next, the degree of dissimilarity of between-language vs. within-language patterns (on-diagonal vs. off-diagonal) was computed using a pairwise *t*-test based on the difference of Fisher-transformed mean correlations ^40^. Significant differences in an area within the searchlight sphere indicated this region differentially encodes L1 and L2. The center of the searchlight was then moved to the next location of the search space, and the statistical analysis was repeated, ultimately yielding a statistical map of all voxels falling within the conjunction map. Analyses were performed using coefficient maps in MNI space. Once single-subject searchlight results were computed, a group statistic was created via a one-sample t-test, which identified voxels showing significantly greater representational similarity within-language than between-languages. Next, we computed random-effect cluster statistics corrected for multiple comparison (*cosmo_montecarlo_cluster_stat*) with a mean of zero under the null hypothesis and 10,000 iterations, and significant clusters were converted to *z*-scores.

**Figure 1.**
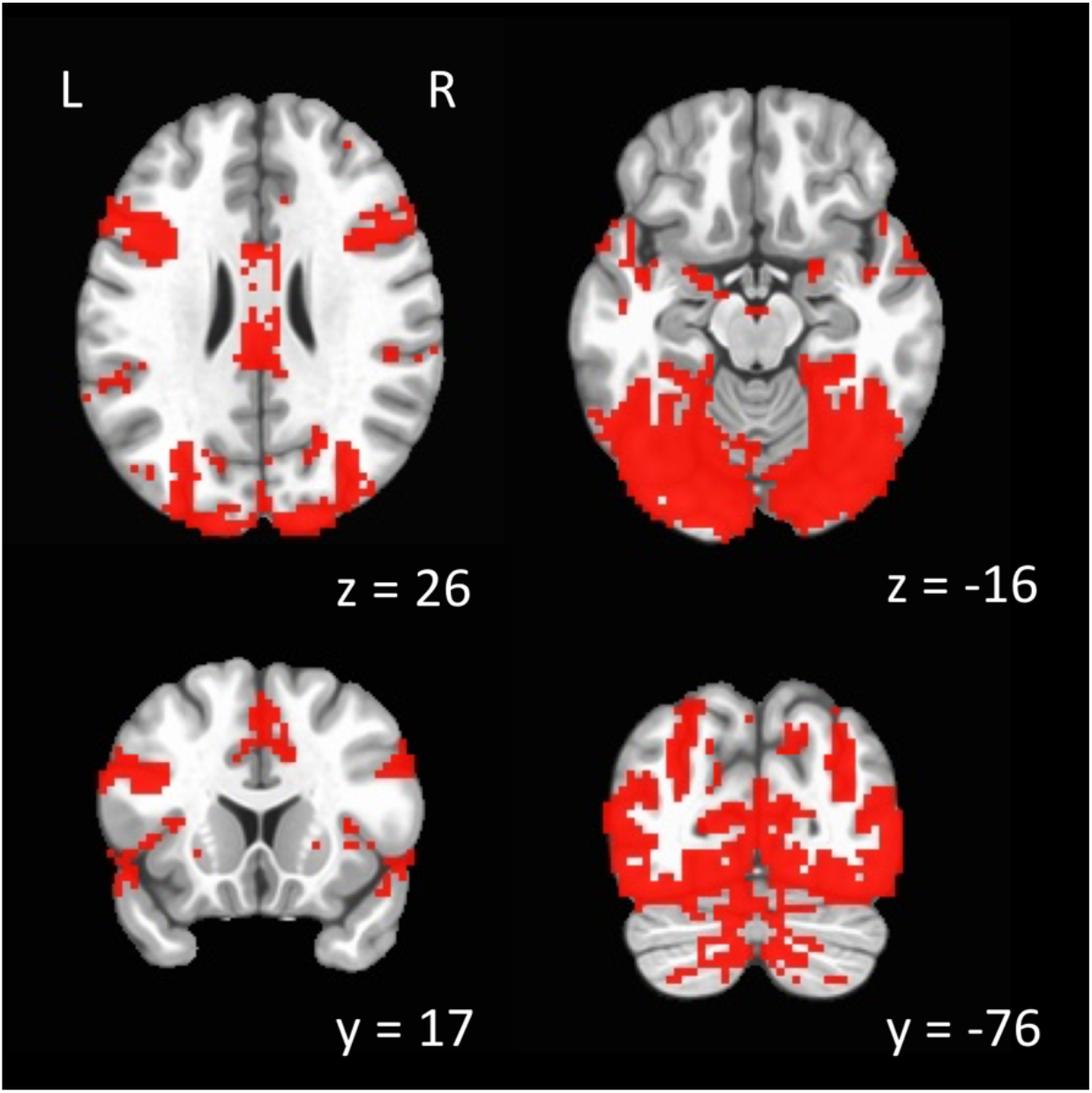
Areas that significantly activated for both L1 English and L2 Mandarin at *p* = .01 uncorrected. Results are overlaid on a stereotaxic brain in MNI space. L=Left, R= Right.

### 2.7 Regression with proficiency and age of acquisition

We then conducted linear regression to examine whether AoA and the difference in proficiency level between L1 and L2 predicted the degree of representational dissimilarity within-subject. Two linear models were constructed, the first with the difference in L2-L1 proficiency as a continuous regressor and adjusting for AoA, the second with AoA as a continuous regressor and adjusting for the difference in L2-L1 proficiency. The minimum cluster-size threshold was determined in two steps. First, we estimated the smoothness of the residuals for each subject output by 3dDeconvolve using the autocorrelation function (ACF) option (AFNI 3dFWHMx), and the mean smoothness level was calculated. Next, minimum cluster size was determined using a 10,000 iteration Monte Carlo simulation (AFNI 3dClustSim) at a voxelwise alpha level of *p* = 0.01, using bi-sided thresholding and first-nearest neighbour clustering. Correction for multiple comparisons at *p* = 0.01 was achieved by setting a minimum cluster size of 7 voxels.

## 3. Results

### 3.1 Behavioural

Performance on the L1 (English) proficiency test ranged from 72.92% to 100%, and performance on the L2 (Mandarin) proficiency test ranged from 12.5% to 77.08%. Analysis of the proficiency test data acquired prior to scanning indicated that L2 proficiency was significantly lower than L1 proficiency (*M* = 88.93%, *SD* = 5.88, *M* = 38.54%, *SD* = 18.15, respectively; *t*(31) = −15.93, *p* < .001, 95% CI [43.94, 56.84]). L2 proficiency did not significantly correlate with L2 AoA (*r*(30) = −0.21, *p* = .255). Participants responded faster on English trials than Mandarin trials (*M* = 1203.60 ms, *SD* = 202.21, *M* = 1607.88 ms, *SD* = 231.78, respectively; *t*(31) = −14.67, *p* < .001, 95% CI [-460.48, −348.09]) and were more accurate on English trials than Mandarin trials (*M* = 94.17%, *SD* = 4.21, *M* = 83.07%, *SD* = 10.35; *t*(31) = 6.84, *p* < .001, 95% CI [7.78, 14.40]).

### 3.2 Conjunction analysis

Results of the conjunction analysis are shown in Figure 1 and Table 2. Both L1 English and L2 Mandarin produced significant activation at a voxelwise *p*-value of 0.01 (uncorrected) in an extensive network of bilateral brain regions including the Heschl’s gyrus, superior temporal gyrus (STG), inferior frontal gyrus (IFG), fusiform and lingual gyri, and occipital and parietal cortices.

### 3.3 Searchlight with split-half correlation analysis

#### 3.3.1 Group-level RSA

As a group, no regions showed significantly greater representational similarity within-language (Mandarin-Mandarin and English-English) compared to between-language (Mandarin-English).

#### 3.3.2 Regression with proficiency and age of acquisition

In order to determine whether proficiency or AoA predicted the degree of difference in representational similarity within- and between-language, subject-wise searchlight maps were submitted to linear regression. L2-L1 Proficiency difference predicted greater within-language representational similarity than between-language similarity in several areas including the left fusiform, IFG, bilateral STG, and right lingual gyrus, shown in Figure 2 and Table 3. All areas showed a positive relationship, indicating that as the difference in proficiency between languages increased, so did the degree of difference in representation between English and Mandarin.

**Figure 2.**
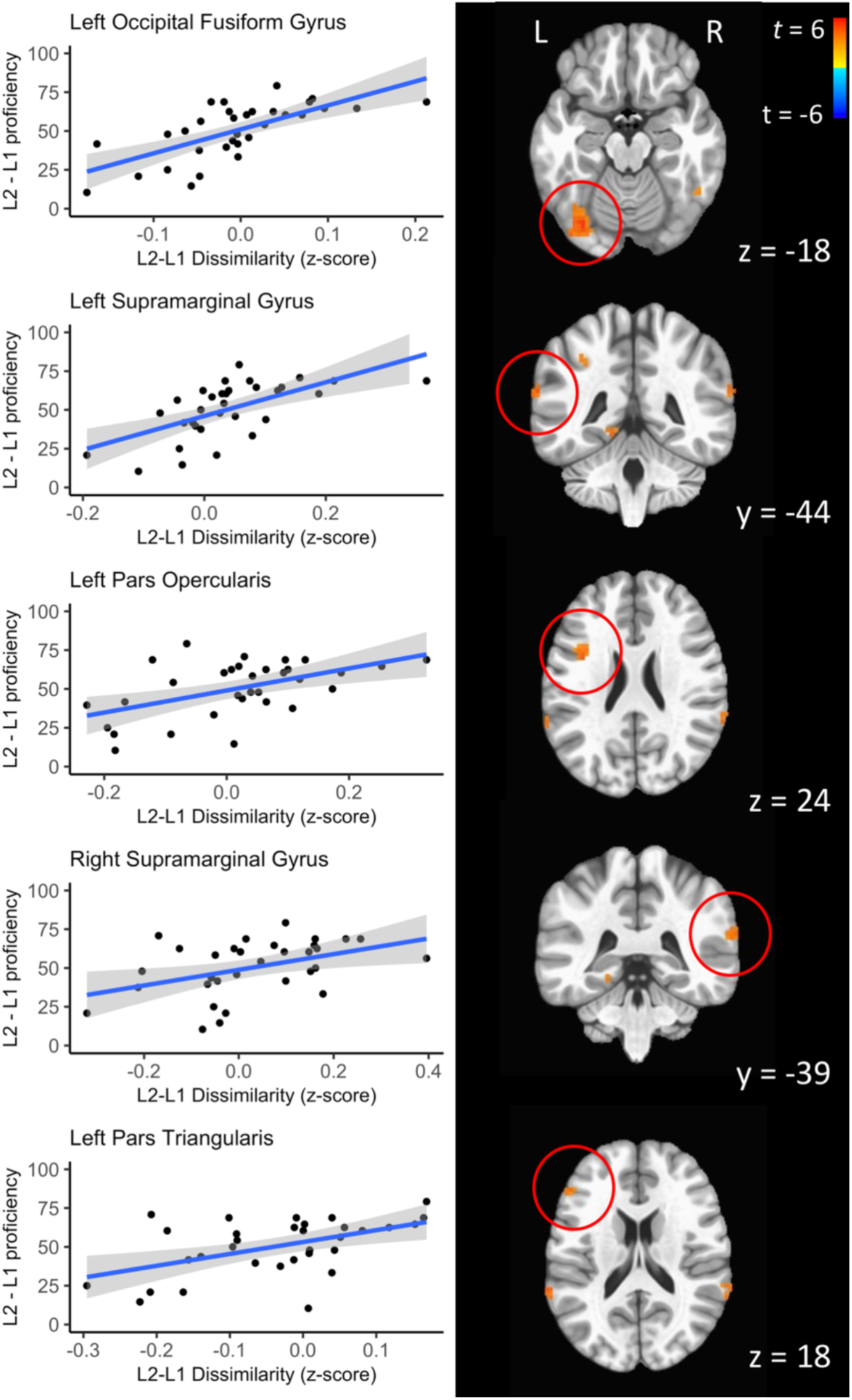
The relationship between difference in L2-L1 proficiency and L2-L1 similarity. *Z*-score values represent the mean across the entire ROI. Higher *z*-scores indicate greater differences between correlation values between-language vs. within-language. Statistical maps are thresholded at *p* = .01, overlaid on an MNI brain atlas. L = left, R = right. Cluster locations and sizes are reported in Table 3.

**Table 3.** Regions where proficiency or AoA significantly predicted *z*-score

|  |  | MNI |  |  | Voxels | *t* | *p* |
| --- | --- | --- | --- | --- | --- | --- | --- |
| Predictor | Region | coordinates |  |  |  |  |  |
|  |  | *x* | *y* | *z* |  |  |  |
| L2-L1 Proficiency | R Cerebellum | 6 | -81 | -30 | 35 | 5.23 | < .001 |
|  | L Occipital fusiform gyrus | -33 | -75 | -18 | 96 | 4.72 | < .001 |
|  | L Supramarginal gyrus | -66 | -45 | 21 | 13 | 4.2 | < .001 |
|  | L Precentral gyrus | -33 | -9 | 66 | 12 | 4.1 | < .001 |
|  | L Pars opercularis | -39 | 3 | 24 | 20 | 4.07 | < .001 |
|  | R Cerebellum | 33 | -66 | -48 | 25 | 4.03 | < .001 |
|  | R Middle occipital gyrus | 51 | -81 | 0 | 9 | 3.97 | < .001 |
|  | L Anterior intra-parietal sulcus | -36 | -48 | 42 | 9 | 3.9 | .001 |
|  | R Primary visual cortex | 18 | -60 | 9 | 26 | 3.88 | .001 |
|  | R Inferior temporal gyrus | 48 | -51 | -24 | 12 | 3.76 | .001 |
|  | L Lingual gyrus | -15 | -45 | -9 | 13 | 3.71 | .001 |
|  | R Supramarginal gyrus | 66 | -45 | 24 | 17 | 3.7 | .001 |
|  | R Orbitofrontal cortex | 33 | 33 | -3 | 10 | 3.63 | .001 |
|  | L Pars triangularis | -51 | 33 | 18 | 8 | 3.61 | .001 |
|  | R Visual cortex ventral V3 | 21 | -78 | -6 | 8 | 3.51 | .001 |
|  | R Cerebellum | 3 | -57 | -45 | 7 | 3.31 | .003 |
| AoA | R Visual cortex ventral V3 | 42 | -93 | -6 | 8 | 5.09 | < .001 |
|  | L Middle temporal gyrus | -54 | -24 | -9 | 9 | 4.34 | < .001 |
|  | L Anterior intra-parietal sulcus | -39 | -45 | 48 | 15 | -3.5 | .002 |
|  | R Insula | 39 | 21 | 0 | 16 | -3.46 | .002 |
|  | R Calcarine sulcus | 18 | -51 | 9 | 8 | -3.32 | .002 |

Note. Coordinates denote the location of peak activation. L/R = Left/Right.

The relationship between AoA and representational similarity is shown in Figure 3 and Table 3. AoA positively predicted greater within-language than between-language representational similarity in the left middle temporal gyrus and right inferior occipital gyrus, indicating that later AoAs were associated with larger differences between L1 and L2 in these areas. In contrast, AoA showed a negative correlation with the left inferior parietal lobe and right insula and calcarine sulcus, indicating that earlier AoAs were associated with smaller L1-L2 representational differences in these areas.

**Figure 3.**
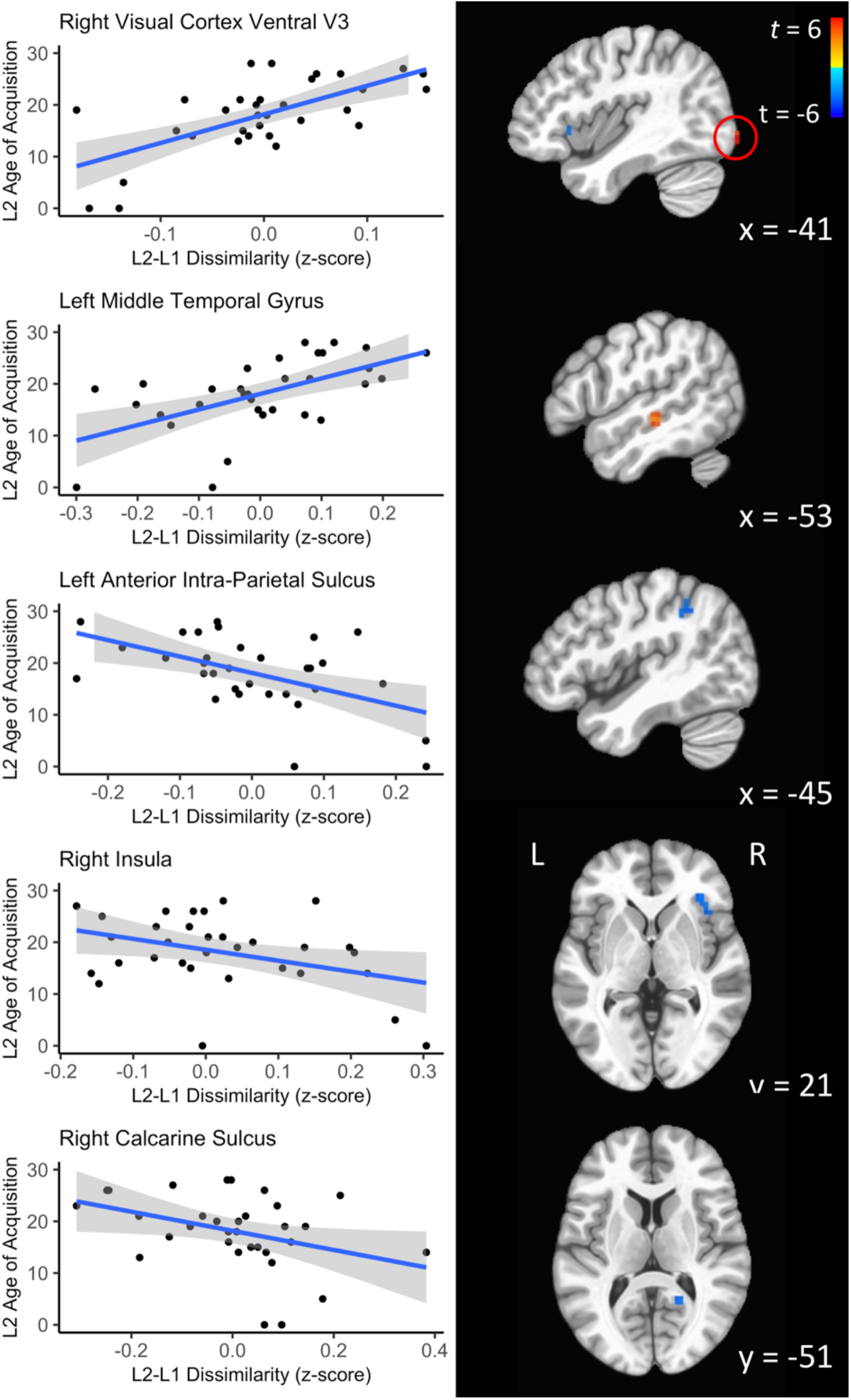
The relationship between L2 AoA and L2-L1 dissimilarity. *Z*-score values represent the mean across the entire ROI. Higher *z*-scores indicate greater differences between correlation values between-language vs. within-language. Statistical maps are thresholded at *p* = .01, overlaid on an MNI brain atlas. L = left, R = right. Cluster locations and sizes are reported in Table 3.

## 4. Discussion

The present study investigated the hypothesis that bilinguals maintain similar, overlapping lexical representations for both of their languages. Using a lexico-semantic recognition task, we found both similarity and dissimilarity in the representation of bilinguals’ two languages within the bilingual word recognition network. There were no regions that significantly differed in their representation of English and Mandarin at the group level, however both proficiency and AoA predicted the degree of representational similarity in several areas. That is, individual differences predicted differentiation in the representation of bilinguals’ two languages in areas that were significantly activated during the word recognition task in both Mandarin and English. These results extend behavioral and ERP findings that bilinguals have a single, integrated lexicon ^12,41–43^, demonstrating how the neural representations within activated regions change with language experience. While prior meta-analyses and reviews have argued this on the basis of relative intensity of fMRI activity ^6,12^, degree of activation cannot tell us about how each language is being represented.

Consistent with our hypotheses, several regions of the language network showed patterns of representation that differentiated languages depending on individual differences. For example, one of these regions was the left IFG (including both the pars opercularis and the pars triangularis), an area engaged in representing and planning articulatory codes for speech and tone ^44–47^. Indeed, these features differ between English and Mandarin in that each language has phonological features that are not present in the other (e.g., tone in Mandarin, consonant clusters in English). The left IFG showed greater representational similarity between languages when the difference between L1 and L2 proficiency was smaller, suggesting that as bilinguals become more matched in proficiency across their two languages, the phonological representations become more integrated. Similarly, language similarity within the bilateral supramarginal gyrus was greater with smaller proficiency differences, an area important for auditory-motor integration during word recognition ^48^.

One notable result was that of representational dissimilarity in lower L2 proficiency and later AoA speakers throughout the ventral visual stream, a cortical pathway responsible for object recognition and concept representation ^49^. The separate representation in visual areas is especially interesting as participants in the present study saw the same images in each language; the manipulation here was only the language in which they heard the names of these objects. As a result, language-dependent differences in this region indicate that this reflects a top-down modulation of high-level visual processing by the linguistic input. Although visual processing of the same images may appear to be a domain-general process, support for it being language-specific comes from the label-feedback hypothesis, which suggests that language modulates ongoing cognitive and perceptual processing ^50^. In line with this hypothesis, each language’s verbal label for the paired image influences the perception of that image. Thus, while the image remains the same, the top-down influence of the language is producing separable representations in high-level visual areas, distinguishing between the visual perception of the spoken word *table* vs. that of the spoken word 桌子 (the Mandarin word for *table*).

There have been numerous studies showing activation differences between L1 and L2, showing greater activation in language areas for one language versus another ^7,51,52^, or showing additional areas recruited for L2 processing vs. L1 processing ^9^. These differences have largely been attributed to later acquisition of L2, differences in proficiency, or other external factors affecting how L2 was acquired ^6,51,53^. In contrast, matched bilinguals tend to show overlapping activity in language regions, with little or no differentiation between languages at the univariate level ^54–56^. L2 speakers in the present study showed experience-dependent representational differences between L1 and L2 in both the language network as well as throughout the ventral visual stream, providing further evidence for integration of bilinguals’ two languages but only when speakers are matched in ability across those two languages.

## Conclusion

We investigated first and second language representation in English-Mandarin bilinguals. Using RSA, we identified both regions in which individual differences predicted differentiation in representation between English and Mandarin. Experience-modulated within-language representational similarity was present in language-network areas (e.g., portions of the left IFG) as well as several regions of the ventral visual pathway, indicating that top-down language knowledge shapes concept representation in the processing stream.

A logical extension of present study is the examination of representational differences in different types of second language processing. For instance, results may differ when comparing two languages that are more similar than English and Mandarin, such as Spanish and French, or when using items that vary in similarity, such as cognates and non-cognates. Additionally, word processing does not involve grammatical processing, which is also an important aspect of bilingual language processing that can differ greatly between L1 and L2. Univariate approaches that contrast degree of brain activation may miss important differences in this regard. The multivariate approach used here may thus provide a way forward in our ability to fully discern how L1 and L2 are represented in the brain.

## Acknowledgments

We are grateful to Suzanne Witt for help with the RSA analysis.

## Author Statement

The authors declare no conflict of interest.

